# Discovery and evaluation of a novel step in the flavonoid biosynthesis pathway regulated by F3H gene using a yeast expression system

**DOI:** 10.1101/753210

**Authors:** Rahmatullah Jan, Sajjad Asaf, Sanjita Paudel, Sangkyu Lee, Kyung-Min Kim

**Affiliations:** Division of Plant Biosciences, School of Applied Biosciences, College of Agriculture & Life Science, Kyungpook National University, 80 Dahak-ro, Buk-gu, Daegu, 41566, Republic of Korea; Natural and Medical Science Research Center, University of Nizwa 616, Oman; College of Pharmacy, Research Institute of Pharmaceutical Sciences, Kyungpook National University, 80 Dahak-ro, Buk-gu, Daegu, 41566, Republic of Korea

**Keywords:** Flavonol 3-hydroxylase, kaempferol, naringenin, nuclear magnetic resonance, yeast episomal plasmid, hydroxylation

## Abstract

Kaempferol and quercetin are the essential plant secondary metabolites that confer huge biological functions in the plant defense system. These metabolites are produced in low quantities in plants, therefore engineering microbial factory is a favorable strategy for the production of these metabolites. In this study, biosynthetic pathways for kaempferol and quercetin were constructed in *Saccharomyces cerevisiae* using naringenin as a substrate. The results elucidated a novel step for the first time in kaempferol and quercetin biosynthesis directly from naringenin catalyzed by flavonol 3-hydroxylase (*F_3_H*). *F_3_H* gene from rice was cloned into pRS42K yeast episomal plasmid (YEP) vector using BamH1 and Xho1 restriction enzymes. We analyzed our target gene activity in engineered and in empty strains. The results were confirmed through TLC followed by Western blotting, nuclear magnetic resonance (NMR), and LC-MS. TLC showed positive results on comparing both compounds extracted from the engineered strain with the standard reference. Western blotting confirmed lack of *Oryza sativa* flavonol 3-hydroxylase (*OsF_3_H*) activity in empty strains while high *OsF_3_H* expression in engineered strains. NMR spectroscopy confirmed only quercetin, while LCMS-MS results revealed that *F_3_H* is responsible for naringenin conversion to both kaempferol and quercetin. These results concluded that rice *F_3_H* catalyzes naringenin metabolism via hydroxylation and synthesizes kaempferol and quercetin.

**Highlights:**

- Current study is a discovery of a novel step in flavonoid biosynthesis pathway of rice plant.
- In this study F3H gene from rice plant was functionally expressed in yeast expression system.
- Results confirmed that, F3H gene is responsible for the canalization of naringenin and converted into kaempferol and quercetin.
- The results were confirmed through, western blotting, TLC, HPLC and NMR analysis.

## Introduction

Flavonoids and isoflavonoids are essential plant aromatic secondary metabolites. They cover a very large family of approximately 9000 known phenolic compounds that mediate diverse biological functions and exert significant ecological impacts. Flavonoids class encompasses approximately 1000 known compounds (Tahara, 2007) including anthocyanin, proanthocyanidin, and phlobaphene pigments, as well as flavonol, flavone, and isoflavone with their respective biological functions in host species (Grotewold, 2005; Lepiniec et al., 2006; Subramanian et al., 2007). With regard to plant biological activities, flavonoids mostly play a role in plant defense system, antimicrobial activity, UV light protection, and fragmentation to different plant parts (Dixon and Steele, 1999; Martens and Mithofer, 2005). Researcher reported that some flavonoids are involved in auxin transport inhibition, have allelopathic activity, and regulate reactive oxygen species in plants (Buer et al., 2010). Among flavonoids, kaempferol and quercetin are the predominant naturally occurring phenolic flavonol compounds with a common flavone nucleus. Both these compounds are of great importance because of their anti-cancer, cardio-protective, and anti-inflammatory activities (Chou et al., 2013). Furthermore, they inhibit the growth of cancer cells, induce apoptosis of cancer cells, and preserve normal cell viability (Chen and Chen, 2013). Kaempferol and quercetin are present in the glycoside form in nature and have numerous biological functions like antioxidant, anti-diabetes, anti-inflammatory, antimicrobial, and anti-fungal activities (Ozcelik et al., 2006; Zang et al., 2011).

In plants, kaempferol and quercetin biosynthetic pathway is well developed and synthesized by complexes of various enzymes, via phenylpropanoid pathway from aromatic amino acids like phenylalanine and tyrosine (Muthukrishnan et al., 2015). At a very early step in the flavonoid biosynthesis pathway, phenylalanine is converted to naringenin through a series of reactions catalyzed by various enzymes. Naringenin is the main intermediate compound responsible for the synthesis of various flavonoids, depending on the enzymes it interacts with (Duan et al., 2017). Similarly, in case of kaempferol and quercetin biosynthesis, flavanone 3-hydroxylase (*F_3_H*) uses naringenin as a substrate and converts it into dihydrokaempferol and further into dihydroquercetin (Deboo et al., 1995). Flavonols (kaempferol & quercetin) are synthesized from dihydrokaempferol and dihydroquercetin when they are catalyzed by flavonol synthase (*FLS*) (Kim et al., 2014). Previous reports showed that kaempferol and quercetin are synthesized by activation of different enzymes in different organisms. For example, functional expression of *Petroselinum crispum*, naringenin 3-dioxygenase (*N3DOX*) in yeast generate dihydrokaempferol which further produces dihydroquercetin kaempferol and quercetin by activation of their respective enzymes (Marín et al., 2018). Similarly, cytochrome P450 flavonoid monooxygenase (*FMO*), which was fused in-frame to the cytochrome P450 reductase (*CPR*) from *Catharanthus roseus*, were expressed in yeast and produced quercetin from kaempferol (Rodriguez et al., 2017). This mechanism showed that kaempferol directly converted to quercetin due to activation of *FMO/CPR*, however in Arabidopsis, dihydrokaempferol first converts into dihydroquercetin due to *F3H* enzyme, which further converts to quercetin by *FLS* enzyme (Yin et al., 2012). In lotus dihydrokaempferol produces kaempferol by regulation of dihydroflavonol reductase (*DFR*) while enzyme responsible for quercetin was unknown (Chen et al., 2013). The proposed pathway of flavonols in *Rubus* shows that, naringenin is hydroxylated by *F3H* being converted in dihydrokaempferol which produced kaempferol by activation of *FLS*. Further kaempferol produce dihydroquercetin which is a precursor of quercetin Gutierrez et al. (2017). Naringenin conversion to different flavonoids is caused by the addition of a hydroxyl group to different positions of the compound, depending on the enzymes interacting with it (Deboo et al., 1995). Previously, Leonard et al. (2006) and Miyahisa et al. (2006) reported kaempferol synthesis in *Escherichia coli* using L-Tyrosin as a substrate and cloned the entire pathway, whereas Leonard et al. (2006) used p-coumaric acid as the substrate of kaempferol and quercetin in *E. coli* and cloned only the downstream pathway. On the bases of proposed pathway, we hypothesized that *F3H* is not only responsible for naringenin conversion to dihydrokaempferol and dihydroquercetin but also responsible for direct conversion to kaempferol and quercetin which is a novel step, not reported up to date. Furthermore it is demonstrated that, *F3H* also converts kaempferol into quercetin.

The aim of the present study is to evaluate the function of *OsF_3_H* gene in the production of kaempferol and quercetin from naringenin. Like other researchers, we did not clone the entire pathway, but cloned only *OsF_3_H* gene the very key step in the flavonoid biosynthesis pathway. To assist with the expression level of the related gene, its function and quantification of quercetin and kaempferol by naringenin as a substrate, in *S. cerevisiae*.

## Material and methods

### Plant material and growth conditions

In the current study, *Oryza sativa japonica* type Nogdong cultivar of Plant Molecular Breeding Lab Kyungpook National University South Korea was used. Before growing, seeds were surface sterilized overnight with 1 ml/l bleach. On the following day, seeds were washed three times with double distilled water and soaked for two days at 34°C. After soaking, the seeds were sown on autoclaved soil and incubated for two days in dark condition. The seeds started to grow in dark condition and were then transferred to a greenhouse, keeping the temperature constant at 25°C, and the RNA was extracted after two weeks.

### cDNA library and PCR

Total RNA was isolated from 2-week-old plant leaves using RNeasy Mini Kit (QIAGEN) Germany, according to the manufacturer’s instructions. Super-standard cDNAs were synthesized using qPCRBio cDNA Synthesis Kit (pcrbiosystem) as per manufacturer’s protocol. To amplify the coding region of *OsF3H* gene, specific primers were designed in NCBI for the PCR reaction. A total sample volume of 50 µl was subjected to the following conditions: initial denaturation at 94°C for 5 min, 40 cycles of denaturation at 94°C for 30 s, annealing at 58°C for 30 s, extension at 72°C for 1 min, and final extension at 72°C for 5 min. The PCR products were extracted using 1% gel and purified using the PCR purification kit.

### Cloning and construction of plasmids

The construct for transformation to yeast was prepared in three steps via restriction-based cloning: (1) insert preparation, (2) construction of vector, and (3) ligation process. To make the insert, the gene was amplified with the forward primer “ggatcc ATGGCGCCGGTGGCCACGACGTT” having the BamH1 site and reverse primer “ctcgag TCACTGCTCTGACGAAGCAACAGAGCAG” with the Xho1 site and purified by gel electrophoresis using Qiagen QIAquick Gel Extraction Kit (Cat # 28706). The purified insert was treated with BamH1 and Xho1 restriction enzymes (New England BioLabs) via incubation at 37°C for 4 h. To prevent methylation, 40 µl of the insert was treated with 2 µl Dpn1 enzyme in the presence of 1 µl Cutsmart buffer (10×) (New England BioLabs) for 2 h at 37°C. At the same time, pRSk42 vector was also treated with BhamH1 and Xho1 enzymes for 4 h at 37°C. After cutting with restriction enzymes, vector was treated with CIP enzyme (New England BioLabs) to reduce the chances of phosphorylation of vector ends. In the last step, the insert was ligated to the vector in ratio of 5:1, respectively, in the presence of Quick Ligase enzyme and 2× Quick Ligase Reaction buffer (New England BioLabs). Before transfer to the yeast, the construct was transformed to *E. coli* JM109 cells via the heat shock method as pRS42k plasmid amplifies in both *E. coli* as well as yeast cells but expresses only in yeast cells. Transformation was confirmed by the isolation of the plasmid DNA from *E. coli*, digestion with restriction enzymes, and the observation of two expected bands of the vector and target gene.

### Strain and media

*E. coli* DH5α and *S. cerevisiae* D452-2 were the strains used in this experiment, having pRS42k plasmid with PGK1 promoter and CYC1 terminator in both *E. coli* and yeast. YPD media (1% yeast extract, 2% peptone, and 2% glucose) were used as the basal media for the routine growth of yeast described by (Sherman, 2002). After autoclaving the solid and liquid media, 150 mg/L of Geneticin (G418) and Spectinomycin (Invitrogen) each were added as selection markers after cooling.

### Transformation to yeast

*S. cerevisiae* transformation was carried out via lithium acetate/single stranded carrier DNA/polyethylene glycol (LiAc/SScarrierDNA/PEG) method (Gietz and Schiestl, 2007). The yeast strain was grown overnight in 10 ml YPD medium at 30°C in a shaker at 200 rpm; further, 250 ml YPD culture flask was incubated. After 12–14 h incubation, the titer of yeast culture was determined by adding 10 µl cells into 1 ml water in a spectrophotometer cuvette and observing the OD at 600 nm. Subsequently, 2.5 × 10^8^ cells were added to 50 ml of pre-warmed YPD into a pre-warmed flask with the titer of 5 × 10^6^. The flask was incubated at 30°C and 200 rpm for about 4 h, after which the cells were harvested and washed with 30 ml water and again rewashed in 1 ml water. Finally, the cells were resuspended in 1 ml of water by vortexing. Simultaneously, single stranded carrier DNA (Salmons perm DNA, Sigma Chemical Co. Ltd, cat. no. D-1626) was incubated for 5 min in a boiling water bath for denaturation and chilled on ice immediately. PEG 50% w/v (add 50 g of PEG 3350 to 30 ml of sterile water) and LiAC 1.0 M (autoclave 10.2 g LiAC with 100 ml water for 20 min) were prepared. Following this, 360 µl transformation mix (35 µl plasmid, 36 µl LiAC, 240 µl PEG, and 50 µl of SS carrier DNA) was added to 100 µl of transformation tube (yeast cells) and vortexed vigorously. The cells were incubated at 42°C for 40 min in a water bath and then harvested by centrifugation for 30 s at 13000 rpm. The supernatant was discarded and the cells were resuspended in 1 ml distilled water by pipetting and 40 µl of cells were plated on each selection media. The plates were incubated for 3 days at 30°C and the transformants were counted.

### Colony PCR

Following the method described by Ling et al. (1995), fresh yeast colonies were used directly for PCR without plasmid purification to confirm gene transformation. Template DNA was prepared by adding a single colony picked using a sterile toothpick to 30 µl of 0.03 M NaOH; the mixture was vortexed vigorously and boiled for 3 min. This was followed by centrifugation at 5000 rpm for 1 min. The supernatant was discarded and 2 µl of the pellet was used as template DNA. This method is highly accurate compared with the old method of adding yeast directly to PCR tubes. All the PCR conditions were the same as those describe previously except the initial denaturation time, which was set at 5 min. PCR products were evaluated on 1% gel and target DNA was visualized using a gel manager.

### Protein isolation and Western blot analysis

The protein was isolated from yeast strains according to the method described by (Chen et al., 2006) with a minor modification. Yeast strain (10 ml) was collected in a 50-ml falcon tube and centrifuged at 5000 rpm for 5 min at 4°C. The supernatant was discarded and the pellet was resuspended in 5 ml TEK buffer solution (50 mM Tris pH 7.5, 2 mM EDTA, and 100 mM KCl) and centrifuged again at 5000 rpm for 5 min. The pellet was resuspended in 5 ml TES buffer solution (50 mM Tris pH 7.5, 2 mM EDTA, 0.8 M sorbitol, 20 mM mercaptoethanol, and 2 mM phenylmethylsulfonyl fluoride) and disrupted by bead beating. Then, 140 mM PEG3350 and 0.2 g/ml NaCl were added to the supernatant and the mixture was incubated on ice for 15 min. After incubation, the samples were centrifuged for 10 min at 10000 rpm and the pellet was resuspended in 100 µl of TEG solution (50 mM Tris pH 7.5, 2 mM EDTA, and 40% glycerol). Protein concentrations were determined by Bradford method (Bradford, 1976); the standard curve shown in Supplementary Fig. 1. The isolated protein (20 µg) was loaded for separation on 12% polyacrylamide gel as described by Laemmli (1970). After separation on polyacrylamide gel, the protein was electro-transferred to nitrocellulose membrane (using transfer apparatus from Bio-Red) and it was suspended in a blocking buffer (50 mM Tris-Cl pH 7.4, 150 mM NaCl, 0.1% Tween 20, and 5% skim milk) for 90 min at room temperature, similar to the study by (Isla et al., 1998). After washing with TBST (50 mM Tris-Cl pH 7.4, 150 mM NaCl, and 0.1% Tween 20) for 40 min, the membrane was incubated in a corresponding primary antibodies in a 1/1200 dilution and polyclonal anti- Mouse IgG antibody from goat (Invitrogen) as a secondary antibody at room temperature. Immunodetection was performed using ECL Western Blotting Detection Reagents (Amersham, Buckinghamshire, UK) and an Image Quant™ LAS 4000 system (Fujifilm, Tokyo, Japan).

### Extraction and thin layer chromatography (TLC) of kaempferol and quercetin

Kaempferol and quercetin were extracted using 100 ml liquid culture grown for 7 days. The crude extracts were isolated using different extraction chemicals, their ratios and extraction time is given in Supplementary Table 1. Crude extracts were then separated into different fractions using a 100-cm-long silica cylinder. The fractions were dried in a rotary evaporator and dissolved in methanol and ethanol for further analysis on TLC. TLC was performed according to the standard method described by (Harborne, 1998) with minor modifications. Approximately 5 µl of 1 mg/ml extracts and the same amount of standard were loaded on the TLC plate (20 × 20 cm thickly coated with 0.4–0.5 nm silica gel) and dried with a drier; the extracts were then allowed to run in their respective mobile phases in the TLC chamber for 25 min. The developed plates were fully dried for 20 min at room temperature and then directly visualized in the TLC viewer under 366 nm UV light. The detected bands were matched with the reference compounds of kaempferol and quercetin. The matching bands were crumb and collected individually and eluted with methanol for further assistance.

## Results and discussion

### Cloning and kaempferol and quercetin pathway design

In recent years, many synthetic tools have been developed with regard to yeast engineering to produce value added secondary metabolites. The proposed pathway for biosynthesis of kaempferol and quercetin is depicted in (Fig. 1). Previous reports showed that *F3H* is a major member of flavonoid biosynthesis enzymes found in all organisms that enhances kaempferol and quercetin biosynthesis (Dixon et al., 1998). Additionally, researchers overexpressed whole genes related to flavonoid pathway in yeast (Trantas et al., 2009). The precursor of flavonoid pathway, p-coumaric acid synthesize by two ways, either it synthesize from tyrosine in tyrosine ammonilyase (TAL), or from phenylalanine in phenylalanine ammonialyase (PAL) pathway (Rodriguez et al., 2017). Further it is demonstrated that, p-coumaric acid, activated by a 4- coumaroyl-CoA ligase (*4CL*) from *P. crispum* while, chalcone synthase (*CHS*) from *P. hybrid,* and chalcone isomerase (*CHI*) from *M. sativa* were used to convert the resulting p-coumaroyl-CoA into naringenin (Rodriguez et al., 2017). Many downstream metabolite synthesis pathways merge from naringenin depending on the activation enzymes. However, depending on the requirement, researchers cloned the entire pathway or a target step of flavonoid pathway, in the microbial factories (Marín et al., 2018).

**Fig. 1.**
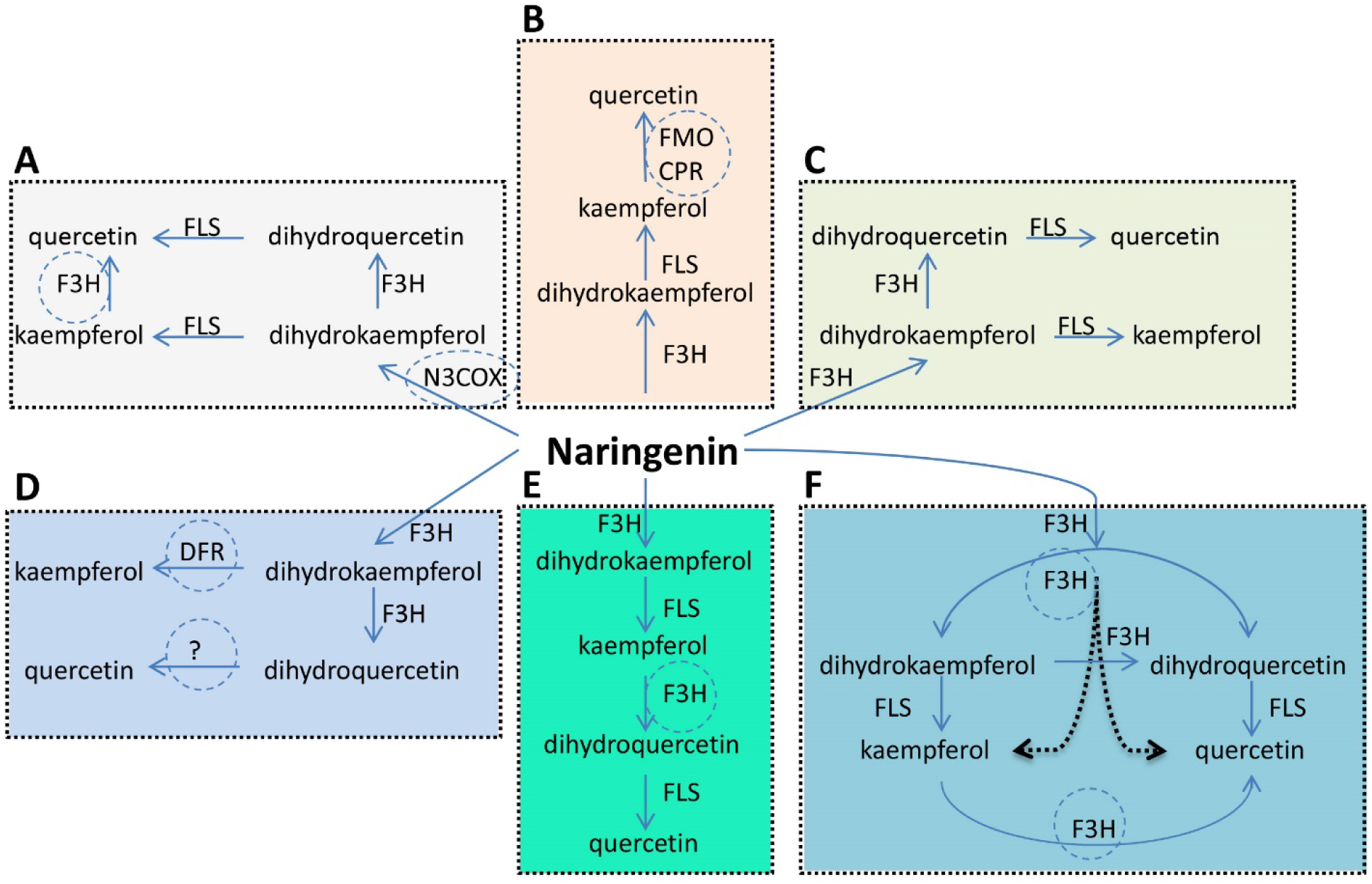
Schematic representation of the flavonoid biosynthetic pathways proposed for the significant production of kaempferol and quercetin in different organisms. Encircled genes shows the differences of gene regulation responsible for different steps regulation among the different organisms pathways. Naringenin is a basic substrate of all these pathways synthesize from phenylalanine which is originated from shikimate pathway through a series of enzymatic reactions. Basically these pathways shows that naringenin converted to dihydrokaempferol by the catalytic activity of F3H and further it also converts dihydrokaempferol into dihydroquercetin. While both of them further converts into kaempferol and quercetin by activation of FLS. (A) In yeast, expression of N3COX is a responsible gene for conversion of naringenin into dihydrokaempferol and also F3H converts kaempferol into quercetin which is different from other organisms. (B) In bacteria, expression of FMO/CPR shows conversion of kaempferol into quercetin. (C) Arabidopsis shows the basic pathway of kaempferol and quercetin synthesis. (D) In lotus, DFR is responsible for conversion of dihydrokaempferol into kaempferol while, gene responsible for quercetin synthesis is unknown. (E) Synthesis of quercetin in blackberry is different from the rest, because kaempferol first converts to dihydroquercetin by activation of F3H which further converts into quercetin by FLS. (E) Our proposed pathway shows that F3H is not only responsible for the conversion of naringenin into dihydrokaempferol and dihydroquercetin, but can also converted naringenin directly to kaempferol and quercetin represented by the dotted arrow. However it also synthesis the intermediate dihydrokaempferol and dihydroquercetin, which then converts to kaempferol and quercetin via FLS activity. We also predicts due to previous study that, F3H is also responsible for the conversion of kaempferol to quercetin.

In this study, we used restriction-based cloning of complete ORF region of *OsF3H* gene ligated into pRS42k expression vector between PGK1 promoter and CYC1 terminator (Supplementary Fig. 2A, 2B). We first transformed the construct to *E. coli* for rapid and efficient multiplication, high copy numbers, and confirmation of ligation. Transformation and ligation were confirmed by plasmid isolation and digestion with corresponding restriction enzymes (Fig. 2A). The same construct isolated from *E. coli* was further transformed to *S. cerevisiae* for functional expression. Approximately 70% of the transformations were successful as 7 colonies were transformed out of 10 and the transformation was confirmed through colony PCR (Fig. 2B) and selection markers. Multiple copy numbers are essential to expresses heterologous genes in yeast; however, significant multiplication of heterologous genes puts pressure on cells and results which may causes uncertainty of the construct. To maintain the stability and optimum copy number of constructs, yeast episomal plasmid (YEp) vector was selected, which replicates autonomously because of the presence of a 2-micron region that acts as origin of replication. The 2-micron origin enhances the copy number, resulting in significant transformation: most of the YEp plasmids range from 5 to 30 copies per single cell (Fang et al., 2011; Mumberg et al., 1995). The engineered strain was induced by subjecting naringenin as a substrate for *OsF_3_H* gene, and the resulting titer was further analyzed using biotechnological tools for confirmation of *OsF_3_H* gene expression, and synthesis of kaempferol and quercetin. Empty and transformed stains of *E. coli* and yeast used in this study are shown in Supplementary Fig S3.

**Fig. 2.**
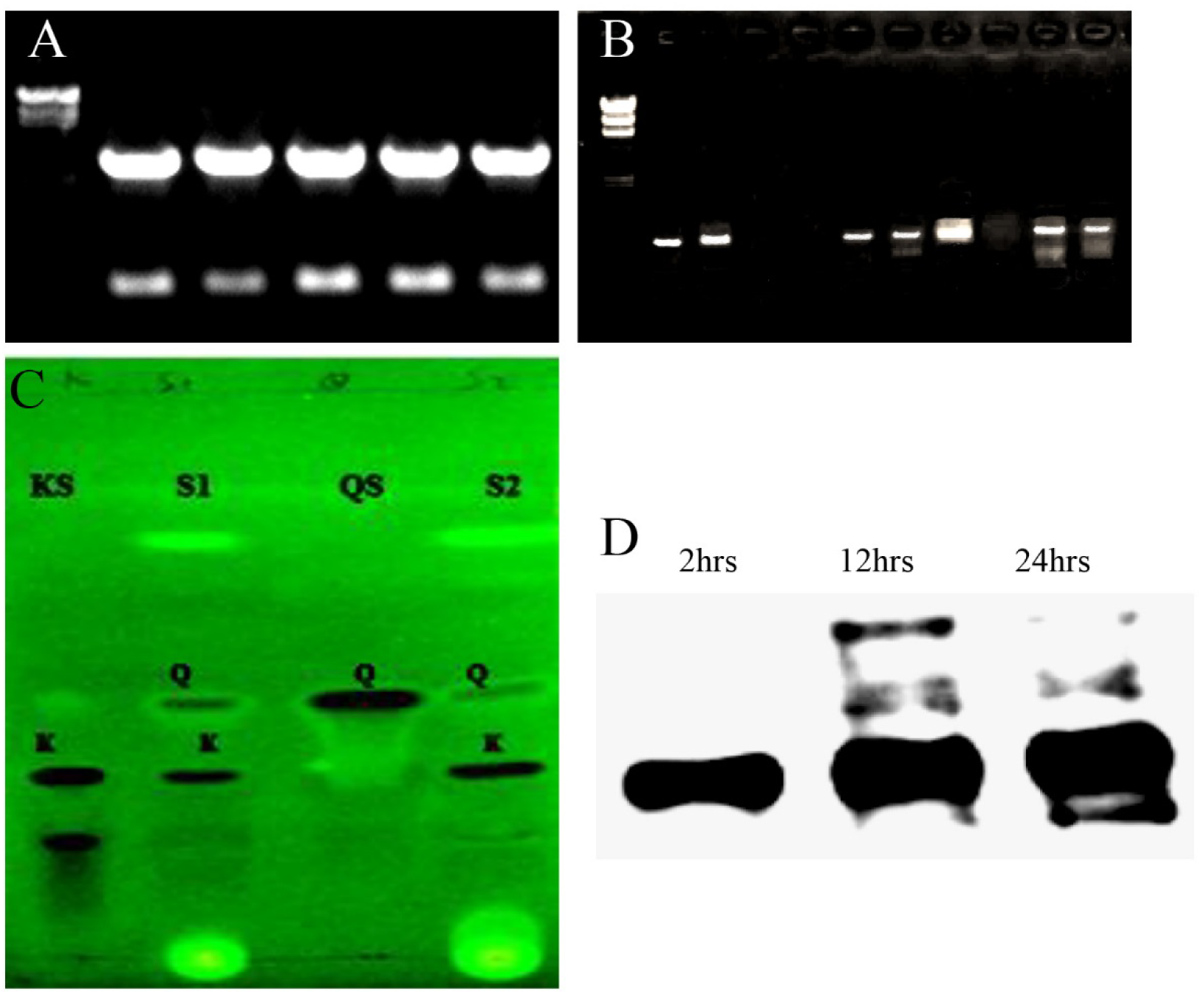
Transformation and functional expression of *OsF3H* gene in yeast. (A) Colony PCR of *OsF_3_H* gene transformed to *S. cerevisiae* (D452-2) strain. (B) Transformation confirmation of *OsF_3_H* gene in *Escherichia coli* cells using pRS42K yeast expression vector. The plasmids of five confirmed colonies are digested with BamH1 and Xho1 restriction enzymes: upper bands show plasmid size, while the lower ones indicate gene size. (C) TLC detection of kaempferol and quercetin at 366 nm UV light. Two samples are analyzed after fractionation through a silica column indicated as S1 and S2, i.e., sample one and two, respectively. At the same time, standards are also run for comparison as represented by KS for kaempferol standard and QS for quercetin standard, whereas K represents kaempferol and Q represents quercetin bands. (D) Immunoblotting of yeast recombinant protein translated from *OsF_3_H* gene. The same amount of transformed strain (each time point) and empty strain protein are prepared and subjected to immunoblotting along with gene-specific antibodies (rabbit anti-CHS chalcone synthase from Sigma Aldrich). SDS-PAGE analysis does not detect the target protein in the empty strains and these are not further analyzed by immunoblotting analysis. At 2, 12, and 24 h represent the time points at which the protein was extracted from yeast strain feeding on substrate.

### TLC analysis

TLC is one of the most important procedures to confirm the presence of related compounds in the extract by comparing the extract with the reference. To substantiate the confirmation of kaempferol and quercetin, TLC was performed using standard method (Harborne, 1998). After fractional distillation and rotary evaporation, a minute quantities (1 mg/ml) of extracts of different profiles were dissolved in their relative solvents along with the same concentration of standard kaempferol and quercetin. TLC profiling was used to indicate the most prominent extract by comparing the extracted kaempferol and quercetin with the standards. Different chromatographic systems (Table 1) of different solvents as a mobile phase were used in random concentration and a favorable solvent that clearly separated kaempferol and quercetin was selected. Different solvent profiles analysis results confirmed that toluene:ethyl acetate:formic acid (7:3:0.5) separated kaempferol and quercetin significantly as shown in Fig. 2C. These results also revealed that there was no difference between Rf of standard and that of the extracted sample. However, the Rf of kaempferol was found higher than that of quercetin in all the mobile phases. Similarly, quercetin was appeared before kaempferol, which shows that quercetin is more polar than kaempferol (Tang et al., 2001). The bands appeared on the TLC plate authenticated that flavonol 3-hydroxylase enzyme catalysed naringenin into kaempferol and quercetin.

**Table 1.** Optimized mobile phases, ratio of solvents used for TLC and Rf values of kaempferol and quercetin

| Chromatographic system | Solvent | Ratio | Rf value of kaempferol | Rf value of quercetin | References |
| --- | --- | --- | --- | --- | --- |
| 1 | ethyl acetate: glacial acetic acid: formic acid: water | 100:11:1<br>1:25 | 0.49 | 0.35 | (Sajeeth et al., 2010) |
| 2 | Benzene: Acetic Acid: Water | 125:72:3 | 0.35 | 0.24 | (Yadav and Kumar, 2012) |
| 3 | n-butanol, acetic acid, water | 4:01:05 | 0.45 | 0.31 | (Yadav and Kumar, 2012) |
| 4 | toluene : ethyl acetate : formic acid | 5:04:01 | 0.36 | 0.24 | (Srivastava et al., 2016) |
| 5 | n-hexane:ethylacetate:acetic acid | 31:14:5 | 0.28 | 0.18 | (Marica et al., 2004) |
| 6 | toluene: ethyl acetate: formic acid | 7: 3: 0.5 | 0.44 | 0.32 | (Kaur and Gupta, 2018) |

### *OsF_3_H* expression in yeast and enzyme assay

To further confirm *F_3_H* activity and protein expression level, the transformed yeast as well as empty yeast (having an empty pRS42K vector) were grown for 24 h at 30°C. After 24 h, naringenin was added to the cultured media of the transformed strain and proteins were isolated at three time points: 2 h, 12 h and 24 h. The enzymatic activity of *F_3_H* was determined by the level of recombinant protein expression among the three time points via Western blotting. Further, immunoblotting was performed with equal protein volumes by making standard curve to estimate the expression level of the recombinant protein. Our results confirmed that the empty vector does not produced the target protein similar to transformed strain which shows lack of *F_3_H* activity. Thus, it was proved that *S. cerevisiae* has no *F_3_H* activity and only transformation with *OsF_3_H* is responsible for the recombinant protein production. On comparing the expression level at each time point, it was assumed that the expression levels slightly increased after each interval because of the continuous catalytic activity of the enzyme (Fig. 2D). This phenomenon predicts that continuous availability of the substrate increases the enzymatic activity of *OsF_3_H* gene because of constant conversion of naringenin into its product. Selection of a suitable and applicable recombinant protein expression system play a key role in protein expression technique. To achieve a high-level protein expression, we selected a well characterized promoter PGK1 and CYC1 terminator, to control the target protein expression of cDNA in host cell, as promoter is the most characterized genetic segment in many yeast expression system (Wagner and Alper, 2016). It has been reported that, heterologous protein production and high degree transferability commonly relied on promotor of *S. cerevisiae* like PGK1 and terminator such as CYC1t (Wagner and Alper, 2016). Selection of a weak promoter can be discouraged due to low level of transcription of foreign gene, and consequently production of low amount of recombinant protein. Similarly, selecting of strong promoter also not recommended all the time due to high level of transcription of foreign gene which can cause a severe stress and can led the cell to death in case of unfolded protein response (UPR). In such case, demands on protein folding and protein size are fundamental for choosing a proper promoter. In this regard we reviewed previous literature for the most frequently using promoter and terminator, and selected PGK1 and CYC1t promoter and terminator respectively and no damage were founded which predicts that the target gene were expressed in a controlled circumstances.

### Identification of kaempferol and quercetin via nuclear magnetic resonance (NMR)

NMR spectroscopy was used to identify a compound by the demonstration of type, number, and position of atoms in a molecule. It is a comprehensive study involving the renovation of the chemical structure of a compound architecture by generating detailed information about carbon and hydrogen atoms in the structure. Currently, NMR application is one of the most significant tools for structure analysis of flavonoids. In the current study, we used NMR spectroscopy to further identify our target compound through structural evaluation. After TLC identification the engineered strain fractions were further analyzed for identification by 1H and 13C NMR (Fig. 3A). Deuteriochloroform is a common solvent used for direct NMR analysis of many flavonoids, generally isoflavones, flavones, and flavonols. We used Methanol D from Cambridge Isotope Laboratories (CIL), USA, as most naturally occurring flavonoids (all flavonoid glycosides) are slightly soluble in deuteriochloroform. However, they are highly soluble in methanol, ethanol, and their derivatives. Samples subjected to NMR analysis revealed only a single compound, the structure was illustrated as quercetin. ^1^H-NMR (600 MHz) δH: 6.41 (1H, d, J = 2.4 Hz), 6.20 (1H, d, J = 2.4 Hz), 7.75 (1H, d, J = 3.0 Hz), 6.91 (1H, d, J = 9.6 Hz), 7.66-7.64 (1H, dd, J = 9.6, 3.0 Hz). 13C-NMR (125 MHz) δC: 148.3 (C-2), 137.5 (C-3), 177.6 (C-4), 162.7 (C-5), 99.5 (C-6),165.9 (C-7), 94.7 (C-8), 158.5 (C-9), 104.8 (C-10), 124.4 (C-1′), 116.3 (C-2′), 146.5 (C-3′), 148.3 (C-4′), 116.5 (C-5′), 121.9 (C-6′). It was recognized as quercetin by comparing with the spectroscopically analyzed data reported in the literature (Ma et al., 2007). With the identification of quercetin and the lack of kaempferol, it is predicted that kaempferol is produced in a very low quantity because NMR detects a very decent amount and highest purity level of compounds. Furthermore, sometimes a single enzyme can catalyze one step of the reaction more efficiently than the other step. It is also possible that the expression of *OsF_3_H* gene in yeast significantly converts naringenin into dihydrokaempferol and dihydroquercetin, and at the same time, also converts the intermediates into quercetin only. The illustrated structure shows the hydroxyl groups attached on 3, 4, 5, and 7 (Fig. 3b) positions.

**Fig. 3.**
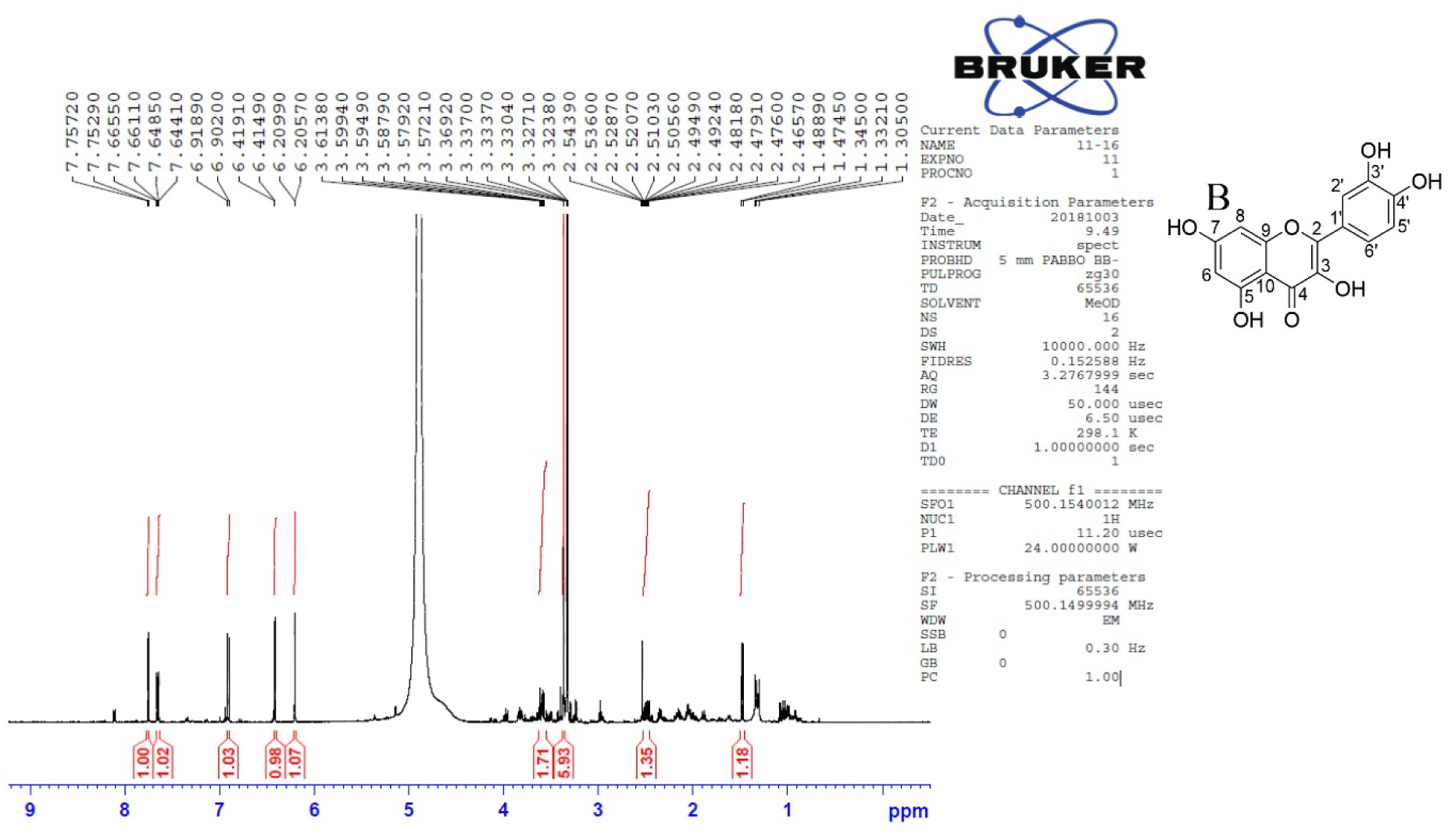
C13 NMR results of quercetin and proposed structure of quercetin. (A) NMR spectra of quercetin, (B) the proposed structure of quercetin.

### *In vivo* activity of *OsF_3_H* in yeast and quantification of kaempferol and quercetin via LCMS-MS

Due to the high pharmaceutical importance of both kaempferol and quercetin, microbial based synthesis of both compounds is a potential approach to enhance its low production in plants. The LC-MS profiles of the *OsF_3_H* assay are shown in Fig. 4. The enzymatic activity of different enzymes expands naringenin metabolism to different pathways responsible for the synthesis of diverse classes of flavonoids. Substrate conversion to kaempferol and quercetin is caused by the addition of single and double hydroxyl groups to naringenin, which is catalyzed by *F_3_H* (Deboo et al., 1995). Our finding shows that using naringenin as a substrate, the recombinant yeast expressing *OsF_3_H* gene successfully metabolized naringenin directly into kaempferol and quercetin which is a novel step discovered and elucidated for the first time. The control strain having an empty vector showed the lack of kaempferol and quercetin accumulation, indicating that *F_3_H* is responsible for naringenin conversion to both compounds. Results showed that metabolism of naringenin increased after each time point, which decreased the naringenin and significantly (p 0.05) increased kaempferol and quercetin (Fig. 4A). LC-MS results also C indicated that *OsF_3_H* metabolized naringenin to kaempferol more significantly than to quercetin. The transformed strain synthesized 8.10 ± 3.81 mg/l, 12.13 ± 5.04 mg/l, and 19.13 ± 1.29 mg/l of kaempferol and 5.42 ± 1.25 mg/l, 7.56 ± 0.86 mg/l, and 8.67 ± 0.39 mg/l of quercetin after 2, 12 and 24 h, respectively. The concentration of naringenin in the titer also decreased gradually along with the increases of both metabolites, showing that *OsF_3_H* strongly catalyzed the reaction (Fig. 4A).

**Fig. 4.**
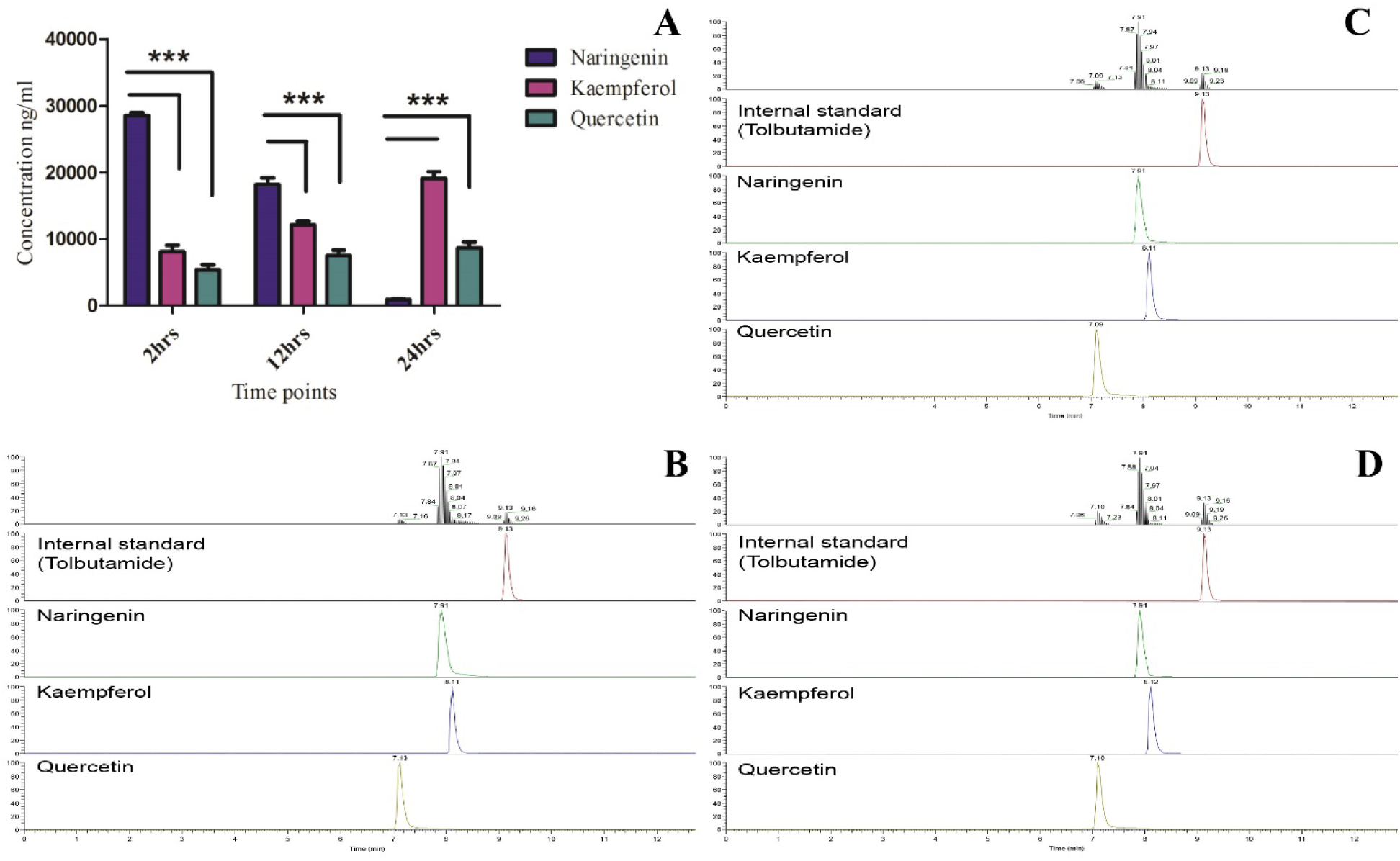
Analysis of naringenin conversion to kaempferol and quercetin using an engineered yeast system expressing *OsF3H* gene using LCMS-MS. (A) LCMS-MS profiling of naringenin, kaempferol, and quercetin. Bars represent mean ± standard deviation, asterisks indicate significant difference (p□0.05 two-way ANOVA, Bonferroni post-test) among naringenin, kaempferol, and quercetin after each time point. (B-D) Extracted ion chromatograms of internal standards (naringenin, kaempferol, and quercetin) in samples extracted after induction in *OxF_3_H* yeast for 2, 12, and 24 h.

Our evaluation and analysis showed that this is a novel finding and *OsF_3_H* gene is responsible for catalyzing naringenin metabolism directly to kaempferol and quercetin without the involvement of *FLS*. Previously it was reported naringenin conversion to dihydrokaempferol via *F_3_H*, which is further converted to kaempferol via flavonol synthase and at last, kaempferol is converted to conversion to quercetin via *F_3_H* (Crozier et al., 2009; Stahlhut et al., 2015) in higher plants. Similarly, Marín et al. (2018) recently reported that naringenin is a substrate metabolized to dihydrokaempferol in the presence of naringenin 3-dioxygenase (*N_3_DOX*), which is further converted to kaempferol and also to dihydroquercetin via *FLS* and *F_3_H*, respectively. Additionally, it has been also reported that, silencing of *FLS* in plants reduces quercetin production which mean that without *FLS* quercetin can also produces (Mahajan et al., 2011). The production of quercetin without *FLS* activity validates that not only *FLS* is responsible for the hydroxylation of dihydroquercetin, which converts to quercetin, but some other enzymes are also responsible for the addition of the OH groups. Flavonoid biosynthesis pathway affirms that naringenin converts to kaempferol and quercetin by hydroxylation depending on enzymes specificity by using O_2_ and NADPH (Leonard et al., 2006). Our finding suggests that *OsF_3_H* gene is responsible for the addition of OH group to naringenin, dihydrokaempferol, and dihydroquercetin for synthesizing both kaempferol and quercetin.

### Conclusion

Using of yeast as a recombinant protein expression system is an easy and short way to characterize a target gene of higher plants. Selection of appropriate plasmid with appropriate promoter and terminator is the key to success of foreign gene expression in the yeast. In current study we used *S. cerevisiae* D452-2 strain and pRS42k plasmid having frequently using PGK1 promoter and CYC1t terminator. Here, we cloned rice *OsF3H* gene in yeast expression vector and functionally expressed in yeast. Here we discovered and evaluated a novel step for the first time in the conversion of naringenin to kaempferol and quercetin by the catalytic activity of *F3H*. The strain containing *OsF3H* gene showed significant synthesis of kaempferol and quercetin by using naringenin as a substrate which is confirmed through TLC, NMR and LCMS-MS analysis while, recombinant protein expression were confirmed through western blotting. With regard to future prospective, this study may help to overexpress *OsF_3_H* gene and assist multifunctional secondary metabolites flavonol biosynthesis in rice plants.

## Supplementary Data

## Acknowledgments

This work was supported by the National Research Foundation of Korea Grant funded by the Korean Government (NRF-2017R1D1A3B04028676), and Brain Korea Plus (BK+). The authors declare no competing interest.

## Author contributions

RJ and KKM designed the experiments. RJ and SA performed experiment and analyzed the data. SP and SL performed HPLC analysis. RJ and SA analyzed NMR data and statistical analysis. RJ and KKM wrote the manuscript.

---

**Fig. S1.**
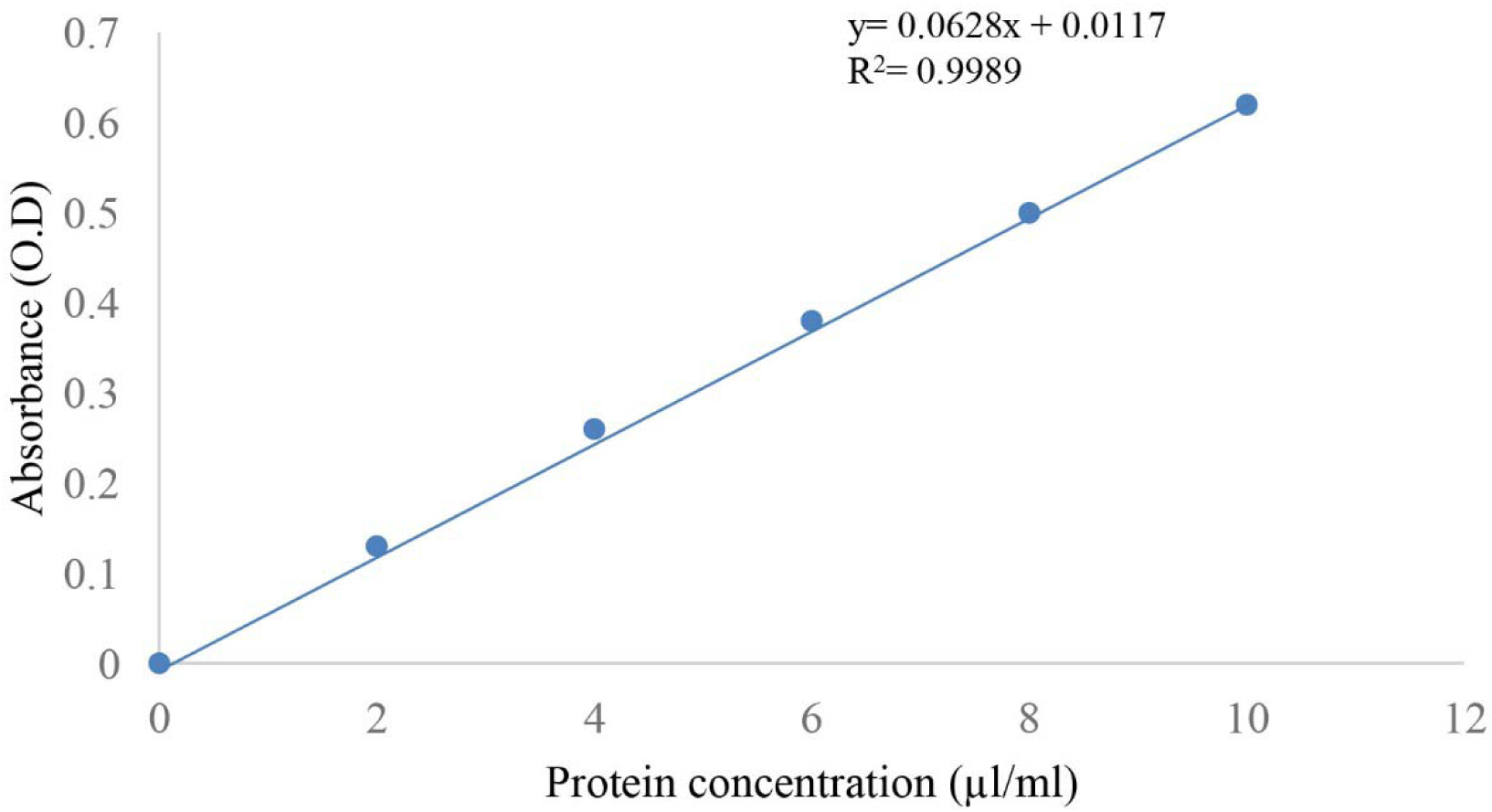
Standard calibration curve for measuring protein concentration using BSA as a protein standard.

**Fig. S2.**
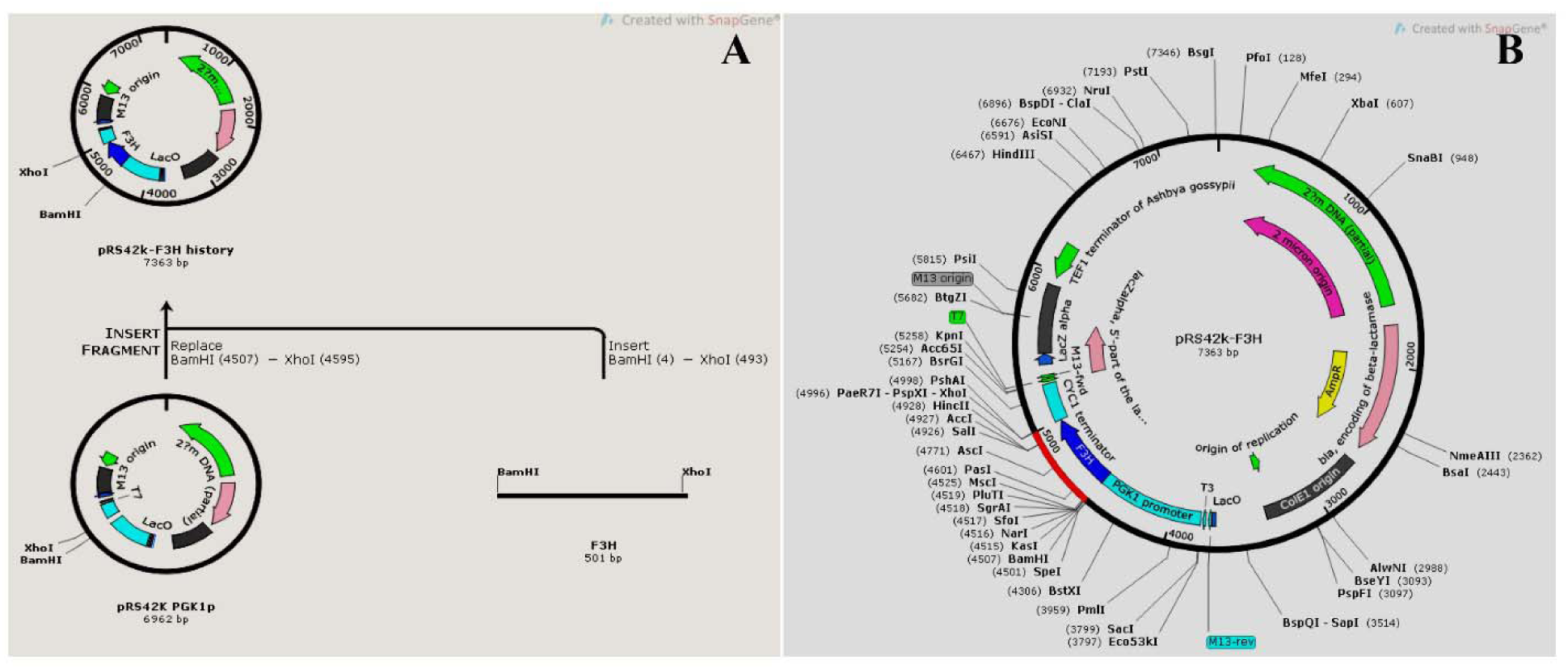
Diagrammatic representation of ligation.

**Fig. S3.**
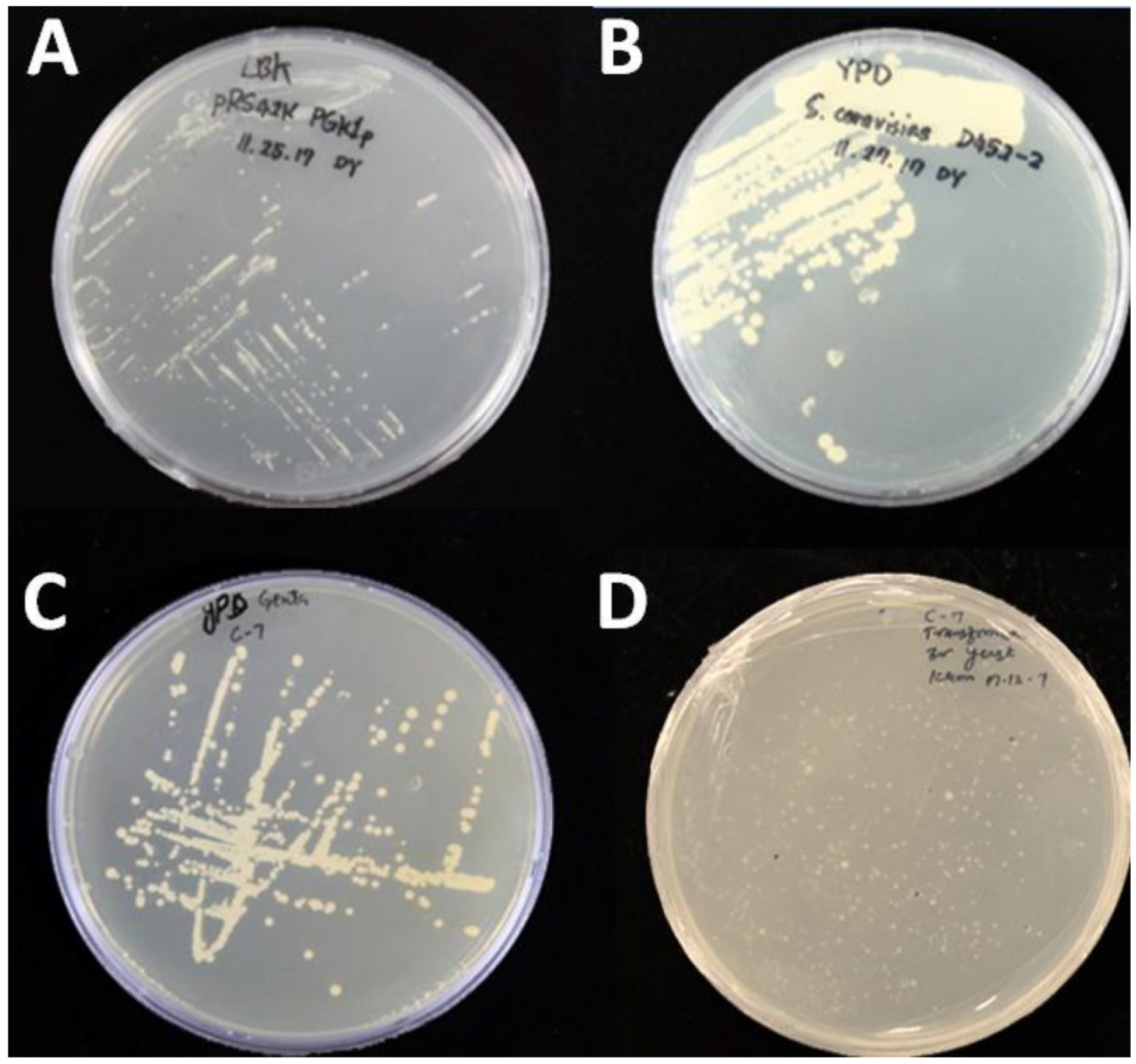
Yeast and *E. coli* strain growth on their respective media.

**Table S1.** Table 1. Solvents and conditions used for kaempferol and quercetin extraction.

| System number | Chemical | Percentage | Condition and time | References |
| --- | --- | --- | --- | --- |
| 1 | Ethanol | 100 | RT/1week | (Batubara et al., 2017) |
| 2 | Methanol | 85 | 3hrs sonication | (Batubara et al., 2017) |
| 3 | Ethanol | 70 | 70°C | (Batubara et al., 2017) |

